# Asymmetric Gait Patterns Alter the Reactive Control of Intersegmental Coordination Patterns during Walking

**DOI:** 10.1101/799213

**Authors:** Chang Liu, James M. Finley

## Abstract

Recovery from perturbations during walking is primarily mediated by reactive control strategies that coordinate multiple body segments to maintain balance. Balance control is often impaired in clinical populations who walk with spatiotemporally asymmetric gait, and, as a result, rehabilitation efforts often seek to reduce asymmetries in these populations. Previous work has demonstrated that the presence of spatiotemporal asymmetries during walking does not impair the control of whole-body dynamics during perturbation recovery. However, it remains to be seen how the neuromotor system adjusts intersegmental coordination patterns to maintain invariant whole-body dynamics. Here, we determined if the neuromotor system generates stereotypical coordination patterns irrespective of the level of asymmetry or if the neuromotor system allows for variance in intersegmental coordination patterns to stabilize whole-body dynamics. Nineteen healthy participants walked on a dual-belt treadmill at a range of step length asymmetries, and they responded to unpredictable, slip-like perturbations. We used principal component analysis of segmental angular momenta to characterize intersegmental coordination patterns before, during, and after imposed perturbations. We found that two principal components were sufficient to explain ~ 95% of the variance in segmental angular momentum during both steading walking and responses to perturbations. Our results also revealed that walking with asymmetric step lengths led to changes in intersegmental coordination patterns during the perturbation and during subsequent recovery steps without affecting whole-body angular momentum. These results suggest that the nervous system allows for variance in segment-level coordination patterns to maintain invariant control of whole-body angular momentum during walking. Future studies exploring how these segmental coordination patterns change in individuals with asymmetries that result from neuromotor impairments can provide further insight into how the healthy and impaired nervous system regulates dynamic balance during walking.

## 1 Introduction

Bipedal locomotion is inherently unstable due to the small base of support, long single-limb support times, and sensorimotor transmission delays [1]. As a result, we must frequently generate corrective responses to maintain balance in response to both internal and external perturbations [2,3]. For example, to recover from unexpected perturbations such as slips or trips while walking, the nervous system generates reactive control strategies involving simultaneous, coordinated responses of both the upper and lower limbs [4,5]. These reactive, interlimb responses to perturbations can restore stability by generating changes in angular momentum that counteract the body’s rotation toward the ground.

One conventional method to capture whole-body rotational dynamics during perturbation responses is to compute whole-body angular momentum (WBAM). WBAM reflects the net influence of all the body segments’ rotation relative to a specified axis, which is commonly taken to project through the body’s center of mass [6–8]. WBAM is highly regulated as its value remains close to zero during normal, unperturbed walking [9,10]. During perturbed walking, angular momentum dramatically deviates from that measured during unperturbed walking [6,7], and this deviation captures the features of body rotation that, if not arrested, would lead to a fall. To regain balance when encountering unexpected perturbations, the central nervous system activates muscles to accelerate body segments and restore angular momentum across multiple recovery steps [11,12].

Angular momentum can also capture balance impairments in populations with gait asymmetries and sensorimotor deficits such as amputees and stroke survivors. These individuals often have a higher peak-to-peak range of angular momentum than healthy controls [13–16], and the presence of gait asymmetries may contribute to balance impairments in these populations. For example, the magnitude of step length asymmetry in people-post stroke is negatively correlated with scores on the Berg Balance Scale, indicating that step length asymmetry is associated with increased fall risk [17].

An important question for clinical researchers is whether there is a causal relationship between gait asymmetry and the ability to maintain balance in response to perturbations during walking. Previous work demonstrated that whole-body dynamics, as measured by WBAM, do not change in response to imposed gait asymmetries in healthy individuals [7]. However, the strategy that the central nervous system uses to stabilize whole-body dynamics remains to be determined. There are two distinct hypotheses capable of explaining the negligible influence of asymmetry on whole-body angular momentum. First, the central nervous system may generate stereotypical, invariant intersegmental coordination patterns in response to perturbations, irrespective of the level of asymmetry. Alternatively, the nervous system could use reactive control strategies that covary with asymmetry in a manner that would lead to invariant control of whole-body momentum. This would be consistent with the uncontrolled manifold (UCM) hypothesis, which predicts that the nervous system allows for variability in segmental angular momenta to stabilize a higher-order performance variable such as whole-body angular momentum [18].

Dimensionality reduction techniques, such as principal component analysis (PCA), are commonly used to capture how the central nervous system coordinates multiple limb segments [6,19]. PCA reduces the high-dimensional, multi-segmental time series data into a lower-dimensional set of latent variables capable of capturing the variance in the overall behavior. Aprigliano et al. used PCA to show that there is no difference in intersegmental coordination patterns between fall-prone older adults and healthy young adults in response to slip-like perturbations [19]. Other studies used PCA of segmental angular momentum to show that the intersegmental coordination patterns observed during recovery from slip-like perturbations are highly correlated with the patterns observed during unperturbed walking [20,21]. Together, these studies suggest that the central nervous system may adopt a preprogrammed and invariant response to perturbation recovery across different tasks and populations.

Here, our objective was to determine how the presence of step length asymmetries influences patterns of intersegmental coordination during slip-like perturbations. Since it has previously been demonstrated that step length asymmetry does not influence the magnitude of whole-body angular momentum, we aimed to determine if this was because the neuromotor system generates stereotypical intersegmental coordination patterns across levels of asymmetry or because the neuromotor system generates patterns of intersegmental coordination that covary with spatiotemporal asymmetry. Ultimately, our findings extend our understanding of how the healthy central nervous system coordinates intersegmental dynamics to maintain balance during walking.

## 2 Methods

### 2.1 Participant characteristics

A total of 19 healthy young individuals (10M, 24 ± 4 yrs old) with no musculoskeletal or gait impairments participated in this study. Lower limb dominance was determined by asking participants which leg they would use to kick a ball. The study was approved by the Institutional Review Board at the University of Southern California, and all participants provided informed consent before participating. All aspects of the study conformed to the principles described in the Declaration of Helsinki.

### 2.2 Experiment protocol

Data used here were collected as part of a previous study [7], and we provide a summary of the procedures and setup below. Participants walked on an instrumented, dual-belt treadmill with force plates underneath (Bertec, USA) at 1.0 m/s for six separate trials and reacted to accelerations of the treadmill belts throughout the experiment. Although 1 m/s was slower than the reported average self-selected speed during treadmill walking [22], we chose this speed to be consistent with other investigations of the role of asymmetry during healthy gait [23–25]. For the first trial, participants walked on the treadmill for three minutes (Baseline) to obtain their natural level of step length asymmetry. Then, for subsequent trials, participants were instructed to modify their step lengths according to visual feedback provided via a display attached to the treadmill, and we informed them that random slip-like perturbations would occur during these trials. The visual feedback displayed the target step length for both right and left legs. A “success” message would appear on the screen if the participants were able to step within three standard deviations of the target step length. Participants completed a randomized sequence of five, six-minute trials with target step length asymmetries (SLA, Eq. 1) of 0%,± 10%,and ± 15% where 0% represents each participant’s baseline SLA.

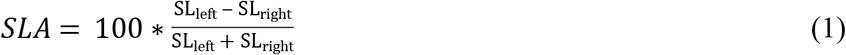

SL_left_ represents left step length and SL_right_ represents the right step length. Each trial consisted of one-minute of practice walking without any perturbations, and then a total of 20 perturbations were applied (10 to each belt) during the remainder of the trial. Foot strike was computed as the point when vertical ground reaction forces reached 150 N. Each perturbation was remotely triggered by preprogrammed Python code and was characterized by a trapezoidal speed profile in which the treadmill accelerated at foot strike to 1.5 m/s at an acceleration of 1.6 m/s^2^, held this speed for 0.3 s, and then decelerated back to 1.0 m/s during the swing phase of the perturbed leg. Participants were aware that they would experience perturbations during the experiment, but the perturbations were randomly triggered to occur within a range of 20 to 30 steps after the previous perturbation to prevent participants from precisely anticipating perturbation timing. This range of steps was also selected to provide participants with sufficient time to reestablish their walking pattern to match with the visual feedback.

### 2.3 Data Acquisition

A ten-camera motion capture system (Qualisys AB, Gothenburg, Sweden) recorded 3D marker kinematics at 100 Hz and ground reaction forces at 1000 Hz. We placed a set of 19 mm spherical markers on anatomical landmarks to create a 13-segment, full-body model [26,27]. We placed marker clusters on the upper arms, forearms, thighs, shanks, and the back of heels. Marker positions were calibrated during a five-second standing trial at the beginning of each trial. We removed all joint markers after the calibration.

### 2.4 Data processing

We post-processed the kinematic and kinetic data in Visual3D (C-Motion, Rockville, MD, USA) and Matlab 2017a (Mathworks, USA) to compute variables of interest. Marker positions and ground reaction forces were low-pass filtered by 4^th^ order Butterworth filters with cutoff frequencies of 6 Hz and 20 Hz, respectively. We selected the type of filter and cut-off frequency based on previous literature [3,28,29]. We calculated the achieved SLA as follows: first, we calculated the mean SLA of the four strides before each perturbation and then distributed these mean values into five equally spaced bins centered at −15%, −10%, 0, 10%, 15% with bin width equal to 5%. We used this achieved SLA instead of target SLA as the independent variable in our statistical analyses. We categorized Baseline (BSL) steps as the two steps before the perturbation occurred, perturbation (PTB) steps as the step during which the perturbation was applied, and recovery (REC) steps as the steps that followed the perturbation. Since we did not find any differences between left and right perturbations, our current analysis includes only perturbations of the right limb [7]. We also focused our analysis on angular momentum about the pitch axis as this was the direction in which the most prominent changes in WBAM were observed. Only minor deviations in WBAM about the roll and yaw axes occurred during the perturbation and recovery steps [7].

### 2.5 Segmental Angular Momentum

We created a 13-segment, whole-body model in Visual3D and calculated the angular momentum of each segment about the body’s center of mass. Segmental angular momenta 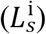 captured how the rotational behavior of each body segment changed in response to the treadmill perturbations. The model included the following segments: head, thorax, pelvis, upper arms, forearms, thighs, shanks, and feet. The limb segments’ mass was modeled based on anthropometric tables [30], and segment geometry was modeled based on the description in Hanavan [31]. All segments were modeled with six degrees of freedom, and we did not define any constraints between segments. Segmental linear and angular velocity were computed using Eq. 2 [15].

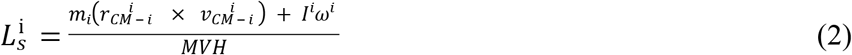

Here, *m*_*i*_ is segmental mass, *r_CM-i_* is a vector from the segment’s COM to the body’s COM, *v_CM-i_* is the velocity of each segment’s COM relative to the body’s COM, *I^i^* is the segmental moment of inertia, ω^*i*^ is segmental angular velocity, and the index *i* corresponds to individual limb segments. Lastly, we normalized momentum by the participant’s mass (*M*), baseline treadmill velocity (*V*), and the participant’s height (*H*) (Eq. 2) following previous literature [9,16]. Since our statistical analysis used a within-subject design, the choice of variables used for normalization should not affect the statistical results.The convention for measuring angular momentum was defined such that positive values represented backward rotation.

### 2.6 Principal component analysis (PCA)

We used principal component analysis (PCA) to extract intersegmental coordination patterns for each step cycle. Before performing PCA, we first time normalized the time series of segmental angular momenta to 100 points for each step cycle. Then, for each participant, we generated an *L*_*s*_ matrix for each achieved SLA (± 15%, ± 10%, ± 5%, %0) and step type (BSL1, BSL2, PTB, REC1, REC2, REC3, REC4) with n_steps*100 rows and 13 columns. On average, we created 6 (achieved SLA) by 7 (step types) matrices per participant as not all participants achieved each desired level of asymmetry. We then standardized each matrix to have zero mean and performed PCA to extract subject-specific coordination patterns using the *pca* function in Matlab’s Statistical and Machine Learning Toolbox. Using PCA, we decomposed the segmental angular momenta data into 1) a weighting coefficient matrix consisting of principal components (PCs) ordered according to their variance accounted for (VAF) and 2) time series scores which represented the activation of each PC throughout the step cycle (Figure 1). We retained the number of PCs necessary to account for at least 90% of variance in *L*_*s*_.

**Figure 1:**
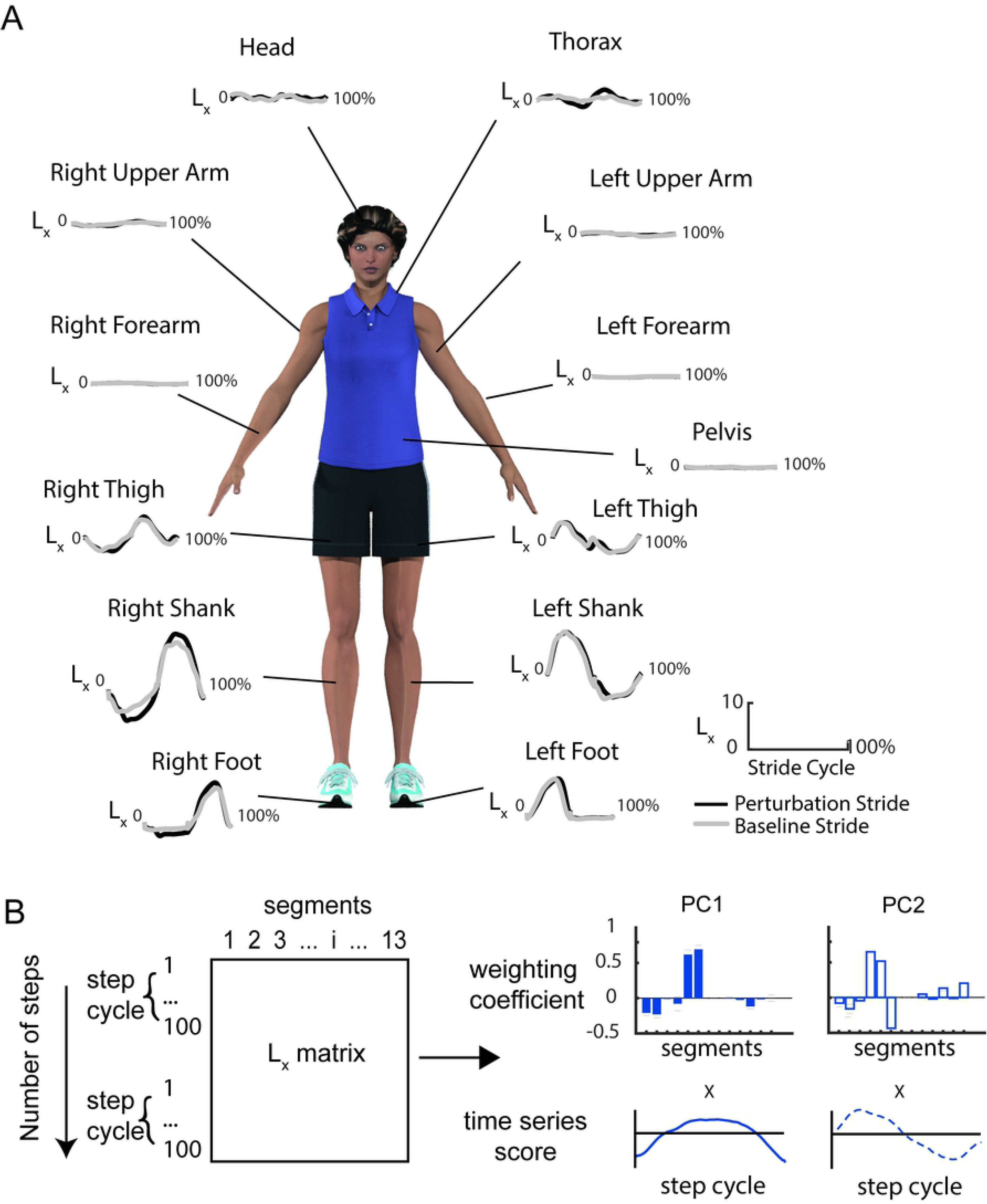
(A) Sagittal plane angular momentum (L_x_) for 13 segments during one representative baseline stride (black) and one perturbation stride (grey). The segments included the thigh, shank, foot, forearm, and upper arm, bilaterally as well as the head, pelvis, and thorax. The duration of each trace is one full stride from 0 to 100% of the stride cycle. (B) Schematic of principal component analysis (PCA) of segmental angular momentum. The organization of the data used as input to the PCA is illustrated to the left. PCA extracts weighting coefficient as intersegmental coordination patterns or principal components (PC1 and PC2) and time series scores of each PC (Filled bar plots: PC1; Open bar plots: PC2).

### 2.7 Comparison of intersegmental coordination patterns

To investigate how intersegmental coordination patterns changed after each perturbation, we compared the PCs extracted from the perturbation and recovery steps to the PCs extracted from baseline steps. We computed the included angle (θ_step_, Eq. 3) between each pair of PCs as this is a common method to compare the similarity between vectors in a high-dimensional space. The included angle of the unit vectors was between 0° (parallel and identical) and 90° (orthogonal and most dissimilar) [32].

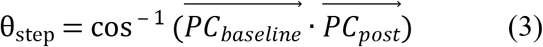

We then determined if the included angle between perturbation steps and baseline steps was outside the distribution of included angles observed during unperturbed baseline walking. To this end, we performed a permutation test that randomly and repeatedly selected two groups of ten baseline steps for each participant. For each permutation, we first performed PCA for each group of 10 steps and then calculated the included angle between the two PCs. We repeated this shuffling process 10000 times for each participant. We used the median of this distribution as a threshold to determine if the included angle for post-perturbation values was greater than what would be expected from step-to-step variance.

Similarly, we computed the included angle between PCs extracted during walking at different levels of asymmetry to those extracted from symmetrical walking to investigate how asymmetry influenced intersegmental coordination patterns. (Eqn. 4).

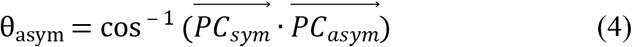

We also determined if the differences in coordination observed during walking with different levels of asymmetry were above the level of variance observed during symmetrical walking. As described above, we obtained a reference distribution of included angles from symmetric walking to determine if the included angle for each level of asymmetry was greater than would be expected from natural, step-to-step variance.

### 2.8 Statistical analysis

All statistical analyses were performed in R (3.4.3) using linear mixed-effects (LME) models. We used the lme4 package to fit the model, the multcomp comparison for multiple comparisons [33], and lmerTest package to calculate p-values [34]. Residual normality was confirmed using the Shapiro-Wilk test. When computing p-values, we used the Satterthwaite approximation for the degrees of freedom based on differences in variance between conditions. We used the Bonferroni correction for multiple comparisons for all post-hoc analyses. For each model, we determined if random effects were necessary by comparing a model including random intercepts for each participant against a model with only fixed effects. The most parsimonious model was chosen based on the results of a likelihood ratio test. The random effects were included to account for the individual differences between subjects. Significance was set at p<0.05 level.

We first determined if the PCs extracted from the recovery steps differed from the PCs extracted from the baseline steps during symmetrical walking. Here, the independent variable was step type, and the dependent variable was θ_step_. The models were fit for both PC1 and PC2. We performed a log transformation of the dependent variable (θ_step_) to ensure that the residuals were normally distributed. Then, we determined if intersegmental coordination patterns during asymmetrical walking differed from those during symmetrical walking. For this analysis, we used Welch’s t-test to evaluate if the included angle between the PCs extracted from the asymmetrical trials and those extracted from symmetric walking were greater than what would be expected by chance. We used Welch’s t-test because the included angle was not normally distributed.

Lastly, we determined if the included angle between each asymmetric trial and symmetric walking varied with the magnitude or direction of asymmetry. For this analysis, the independent variables were the magnitude of asymmetry, the direction of asymmetry, and the interaction between asymmetry magnitude and direction, and the dependent variable was θ_asym_. We fit separate linear mixed-effect models for each of five steps (Baseline1, Baseline2, Perturbation, Recovery 1 and Recovery 2) and each PC. We performed a log transformation of the dependent variable (θ_asym_) to ensure that the residuals were normally distributed.

## 3 Results

For all steps, two principal components accounted for ~95% of the variance in segmental angular momentum (Table 1). On average, PC1 explained 74 ± 4% of the variance, and PC2 explained 22± 1% of the variance, while PC3 accounted for less than 3% of the variance. Thus, the remaining analysis focuses on the first two PCs.

**Table 1:** Variance accounted for (VAF) for PC1, PC2, and PC3 during baseline steps, perturbation steps, and recovery steps.

| Step Type | PC1 | PC2 | PC3 | Sum |
| --- | --- | --- | --- | --- |
| Baseline steps | $74 \pm 4\%$ | $22 \pm 5\%$ | $2 \pm 1\%$ | $98 \pm 1\%$ |
| Perturbation steps | $75 \pm 5\%$ | $20 \pm 5\%$ | $3 \pm 1\%$ | $98 \pm 1\%$ |
| Recovery steps | $74 \pm 3\%$ | $21 \pm 4\%$ | $3 \pm 1\%$ | $98 \pm 1\%$ |
| All steps | $74 \pm 4\%$ | $22 \pm 4\%$ | $2 \pm 1\%$ | $98 \pm 1\%$ |

### 3.1 Patterns of intersegmental coordination when walking with equal step lengths

Contributions from the lower extremities were typically dominant in the first PC, while contributions from the arms, pelvis, thorax, and head were less prominent (Figure 2). During right steps, the left leg was in the swing phase and generated more positive momentum about the body’s COM, while the right leg generated negative momentum. Thus, the weighting coefficients for the left leg segments (left thigh, shank, and foot) were positive while the coefficients for the right leg segments were negative. Similarly, during a left step, the right leg was in the swing phase and generated more positive momentum about COM, while the left leg generated negative momentum. Thus, the weighting coefficients were positive while the coefficients for the left leg segments were negative. Overall, the first PC captured the opposing momenta of the two legs resulting from differences in the direction of rotation relative to the body’s center of mass.

**Figure 2:**
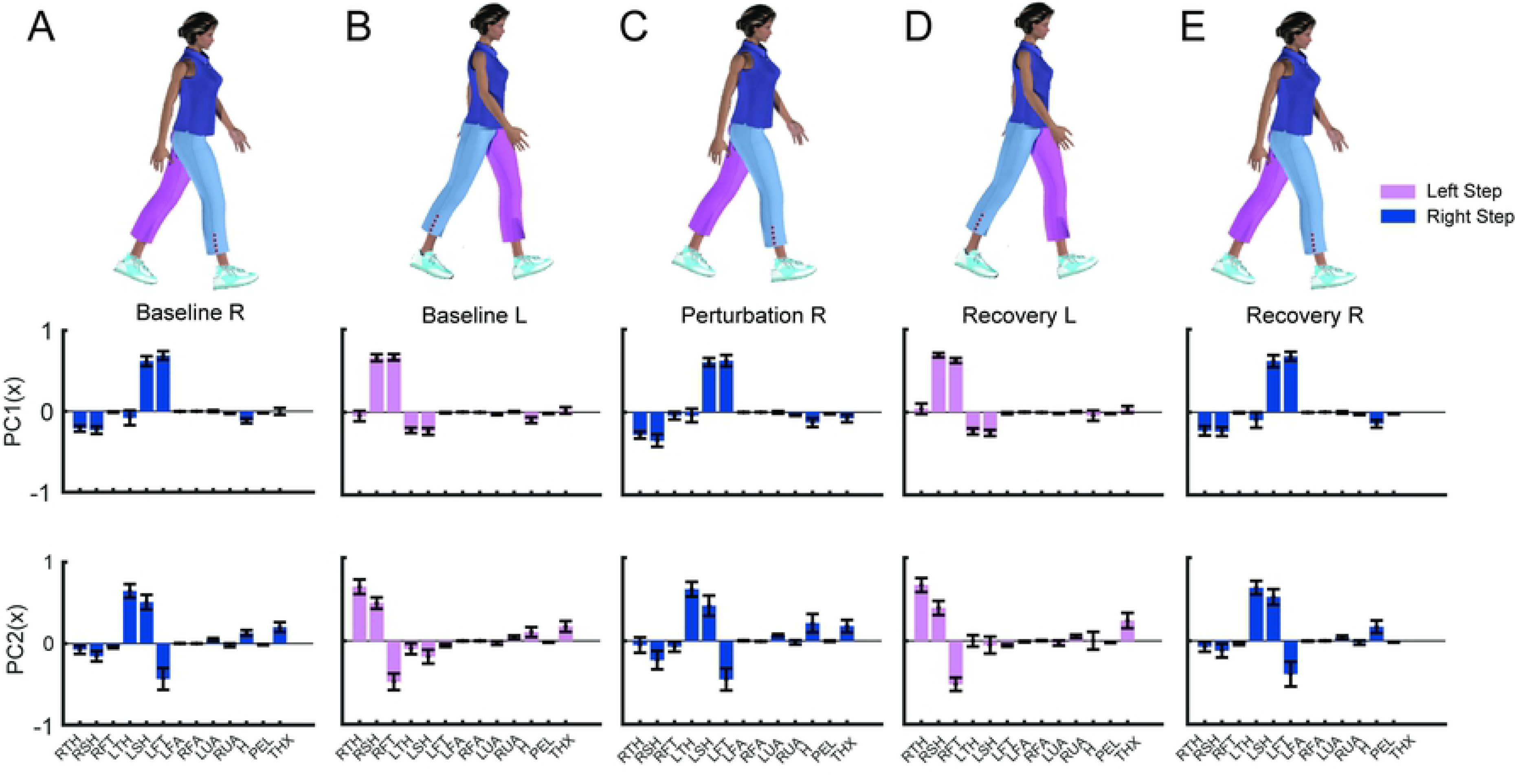
Principal components (PC) extracted from segmental angular momentum during (A) baseline right steps, (B) baseline left steps, (C) perturbation steps, (D) recovery left steps, and (E) recovery right steps when walking symmetrically (N=17). Blue: Right step; Pink: Left step; Filled bars: PC1; Unfilled bars: PC2. The 13 segments include: RTH (right thigh), RSH (right shank), RFT (right foot), LTH (left thigh), LSH (left shank), LFT (left foot), LFA (left forearm), RFA (right forearm), LUA (left upper arm), RUA (right upper arm), H (head), PEL (pelvis), THX (thorax).

For PC2, weighting coefficients for distal segments were also larger than the weighting coefficients for proximal segments, although the coefficient for the thorax (THX) increased compared to that in PC1. During the right step, the left thigh and left shank’s momenta opposed the momentum of the left foot. Similarly, during the left step, the right thigh and shank momenta opposed the right foot momentum. Thus, PC2 captured intralimb cancellation of segmental momenta.

### 3.2 Effects of perturbations on patterns of intersegmental coordination

During the perturbation step, there was a significant increase in the included angle, which indicated that the intersegmental coordination patterns during perturbation steps differed from the coordination patterns during baseline steps (Figure 3). For this analysis, the results of the log-likelihood ratio test revealed that random effects were necessary for the regression model. For PC1, we found that the intersegmental coordination patterns were significantly different from the patterns during baseline walking for the perturbation steps (t(54)=18.2, p<2e-16), first recovery steps (t(54)=11.8, p<2e-16), and second recovery steps (t(54)=8.4, p=2.3e-11). Similarly, for PC2, intersegmental coordination differed during perturbation steps (t(54)=11.8, p<2.0e-16), first recovery steps (t(36)=6.7, p<2e-16),and second recovery steps (t(54)=4.9,p=8.9e-6).There was no significant difference between intersegmental coordination patterns during the third recovery steps for either PC1 (p = 0.97) or PC2 (p = 0.14). Thus, participants generally were able to restore their coordination patterns to baseline by the third recovery step.

**Figure 3:**
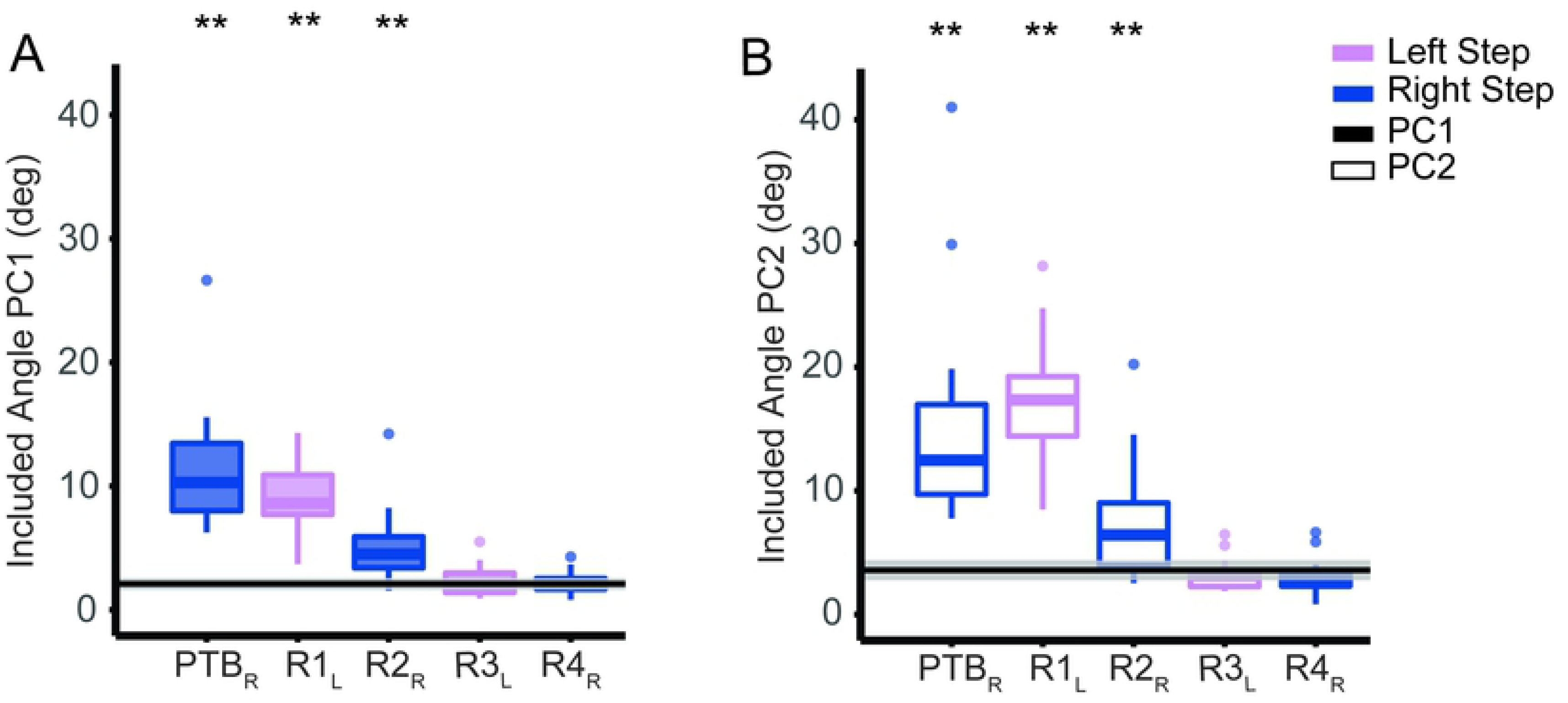
Included angle between PCs extracted during each step relative to baseline steps during symmetric walking (** p<0.001). The horizontal bars and corresponding stars indicate significant differences in the included angle. The data are represented as boxplots such that the lower and upper edges of the box indicate the 25^th^ and 75^th^ percentile of the data, respectively. The horizontal line in each box indicates the median. The whiskers extend to the furthest data point beyond the lower or upper edges of the box that is within a distance of 1.5 times the middle 50^th^ percentile of the data. Dots that lie beyond the whiskers indicate outliers. Blue: Right step; Pink: Left step; Filled box plots: PC1; Non-filled box plots: PC2. The black line indicates the mean of the permutated angle distribution of baseline steps and the shading indicates the standard deviation.

### 3.3 Effects of step length asymmetry on patterns of intersegmental coordination

Although the general patterns of intersegmental coordination were similar across levels of asymmetry, asymmetric walking patterns led to measurable changes in the contributions of the distal lower extremity segments (Figure 4). Qualitatively, we observed increased weights at the left foot as well as decreased weights of the left shank segment for the first principal component during right steps. This likely reflected the need for longer left steps and faster foot swing for positive step length asymmetries.

**Figure 4:**
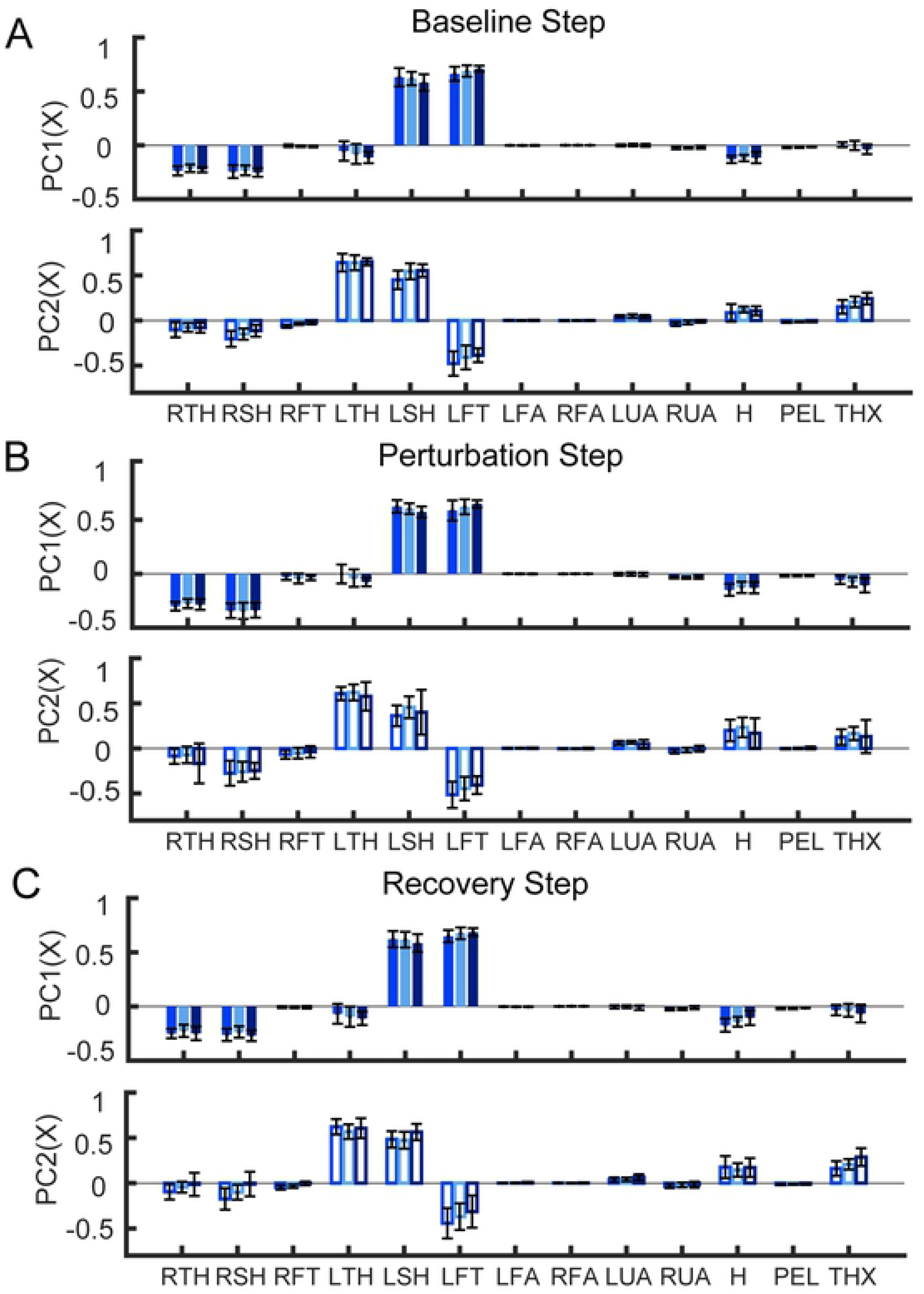
The first intersegmental coordination pattern (PC1) and the second coordination pattern (PC2) during (A) baseline right step, (B) perturbation step, and (C) the second recovery step with −15%, 0% and 15% step length asymmetry. The colored bars indicate the mean value across all participants (N=17), and the black lines indicate the standard deviation.

As the magnitude of achieved asymmetry increased, we observed an increase in the deviation of intersegmental coordination patterns from symmetrical walking (Figure 5). Results of log-likelihood ratio tests showed that random intercepts were required in the regression models. One outlier was removed before fitting the linear mixed model for the perturbation step for PC2 because it was more than three standard deviations higher than the median of the included angles. Excluding the outlier did not change the statistical outcome. All included angles differed from the permutated estimate of included angles (p<0.05), indicating that intersegmental coordination at each level of asymmetry differed from the coordination pattern during symmetrical walking. For all steps, we observed a significant main effect of asymmetry on the included angle between the PCs from the asymmetric trials and the symmetric trial (Table 2).

**Table 2.** Statistical results from the ANOVA examining the effects of asymmetry and direction on the included angle for each step type.

| Step Type | PC | Factor | numDF | denDF | F-value | P value |
| --- | --- | --- | --- | --- | --- | --- |
| Baseline1 | PC1 | **Asym** | **2** | **73** | **9.7** | **<0.001** |
|  |  | Direction | 1 | 77 | 2.4 | 0.13 |
|  |  | Asym:Direction | 2 | 74 | 0.6 | 0.55 |
|  | PC2 | **Asym** | **2** | **72** | **14.4** | **<0.001** |
|  |  | **Direction** | **1** | **78** | **4.0** | **0.049** |
|  |  | Asym:Direction | 2 | 74 | 0.08 | 0.92 |
| Baseline2 | PC1 | **Asym** | **2** | **72** | **5.7** | **0.005** |
|  |  | Direction | 1 | 75 | 1.3 | 0.26 |
|  |  | Asym:Direction | 2 | 73 | 0.1 | 0.88 |
|  | PC2 | **Asym** | **2** | **71** | **11.0** | **<0.001** |
|  |  | Direction | 1 | 74 | 0.007 | 0.93 |
|  |  | Asym:Direction | 2 | 72 | 2.2 | 0.12 |
| Perturbation | PC1 | **Asym** | **2** | **73** | **19.0** | **<0.001** |
|  |  | Direction | 1 | 75 | 1.9 | 0.18 |
|  |  | Asym:Direction | 2 | 73 | 0.5 | 0.59 |
|  | PC2 | **Asym** | **2** | **72** | **8.7** | **<0.001** |
|  |  | Direction | 1 | 73 | 1.3 | 0.25 |
|  |  | Asym:Direction | 2 | 72 | 0.68 | 0.51 |
| Recovery1 | PC1 | **Asym** | **2** | **74** | **11.2** | **<0.001** |
|  |  | Direction | 1 | 78 | 0.1 | 0.74 |
|  |  | Asym:Direction | 2 | 75 | 1.3 | 0.29 |
|  | PC2 | **Asym** | **2** | **72** | **9.1** | **<0.001** |
|  |  | Direction | 1 | 75 | 0.1 | 0.72 |
|  |  | Asym:Direction | 2 | 73 | 0.8 | 0.45 |
| Recovery2 | PC1 | **Asym** | **2** | **72** | **8.7** | **<0.001** |
|  |  | Direction | 1 | 75 | 1.8 | 0.18 |
|  |  | Asym:Direction | 2 | 73 | 0.4 | 0.67 |
|  | PC2 | **Asym** | **2** | **73** | **8.5** | **<0.001** |
|  |  | Direction | 1 | 77 | 1.9 | 0.18 |
|  |  | Asym:Direction | 2 | 74 | 0.2 | 0.84 |

**Figure 5:**
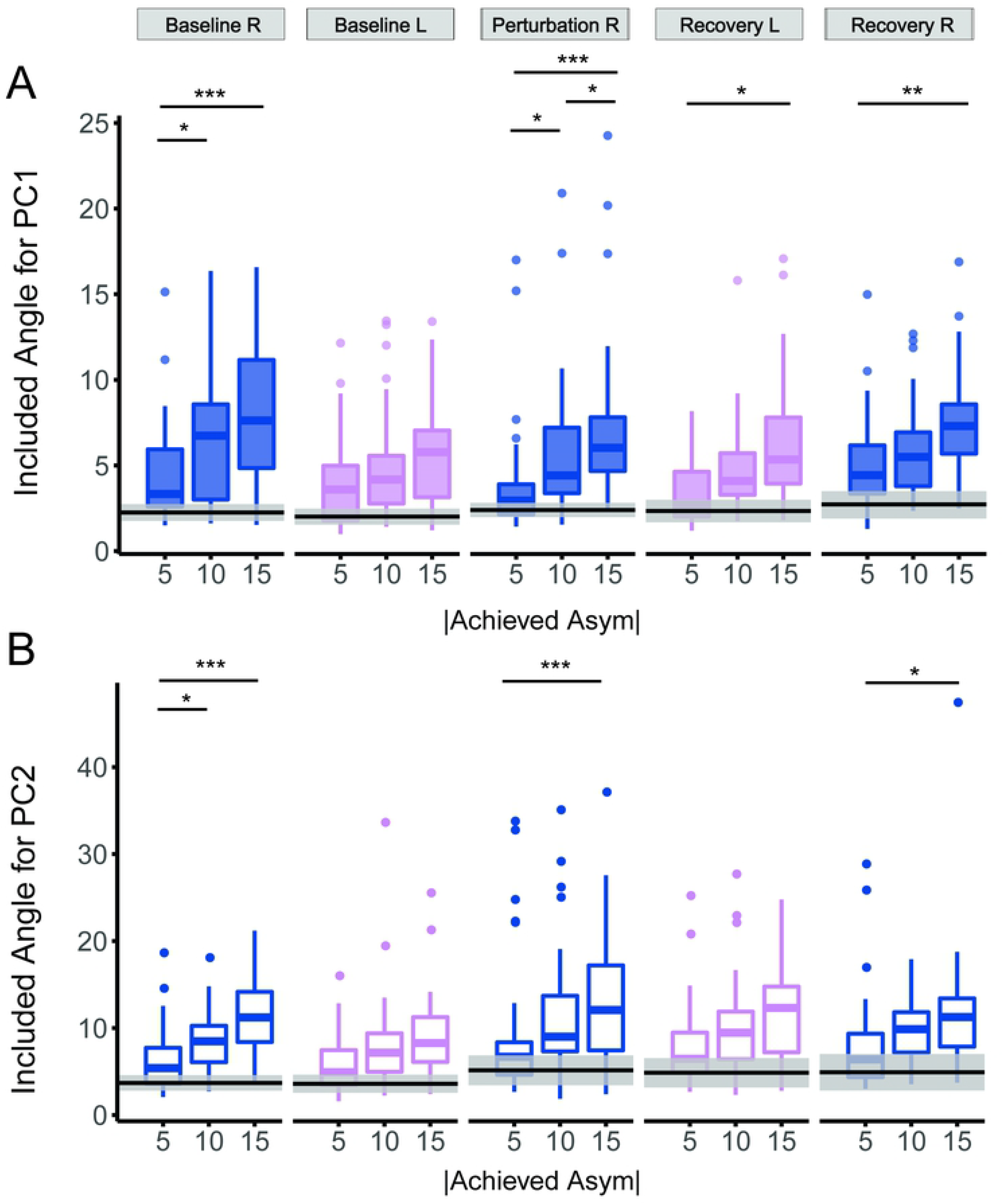
included angle between PCs extracted during asymmetrical walking (5%, 10%, and 15%) and symmetrical walking for each step (*** p<0.001, ** p<0.01, * p<0.05). Blue: Right step; Pink: Left step; Filled box plots: PC1; Non-filled box plots: PC2. The shaded gray area indicated the standard deviation of permutated included angle for each step, and the black line indicated the mean of the distribution.

The included angle between the PCs extracted during asymmetric walking and symmetric walking increased with the magnitude of achieved asymmetry (Figure 5). Specifically, the difference between intersegmental coordination patterns was greater when walking with 15% asymmetry compared to 5% asymmetry during right baseline steps (Bonferroni corrected p<0.001), perturbation steps (Bonferroni corrected p<0.001), first recovery steps (Bonferroni corrected p=0.03) and second recovery steps (Bonferroni corrected p=0.002) for PC1. The difference in included angles was also significantly different from 5% asymmetry for PC2 when walking with 15% asymmetry during baseline right steps (Bonferroni corrected p=0.01) and perturbation steps (Bonferroni corrected p = 0.003) and second recovery steps (Bonferroni corrected p=0.04). Lastly, there was only an effect of the direction of asymmetry for PC2 (F(1, 79), p=0.049) during the baseline right step (Baseline 1).

## 4 Discussion

We investigated how step length asymmetry affected intersegmental coordination patterns during responses to treadmill-based slip perturbations during walking. Our central finding was that intersegmental coordination patterns observed during asymmetrical walking differed from symmetrical walking during both unperturbed walking and perturbation recovery. When combined with previous observations that the reactive control of overall WBAM is not influenced by asymmetry [7], these results indicate that healthy people use a flexible combination of intersegmental coordination patterns rather than invariant reactions to maintain WBAM during perturbation responses when walking with asymmetric gait patterns.

Variations in coordination patterns during asymmetrical walking likely resulted from changes in the momentum generated by the lower extremities to reach the target asymmetry. Since the distal segments of the lower limbs are relatively far from the body’s center of mass and have a high velocity, they make the largest contribution to changes in intersegmental coordination patterns. For example, to achieve a positive asymmetry, participants placed their left foot further in front of the center of mass and increased the extension of their right hip so that the right foot was further behind their COM at heel strike. To achieve this objective, participants had to increase swing velocity. This likely explains why we observed increased weights of the left foot as SLA increased during right steps in the first principal component since positive step length asymmetries required longer left steps and faster foot swing.

The observation that reactive control of WBAM is consistent across levels of asymmetry [7] despite the variation in intersegmental coordination observed here may indicate that WBAM is a task-level variable that is stabilized by the nervous system during perturbation recovery. This is consistent with the framework proposed by the uncontrolled manifold hypothesis (UCM), which argues that the central nervous system allows for variability over a manifold of solutions that all successfully stabilize a higher-level performance variable [35]. Here, WBAM would serve as a high-level performance variable that is stabilized through covariation of elemental, segmental-level momenta. For example, Papi et al. demonstrated a similar concept when they found no differences between people post-stroke and healthy individuals in COM displacement during the stance phase of walking despite between-group differences in lower extremity joint kinematics [36]. Therefore, it is possible that when dynamic stability is challenged during walking, the central nervous system carefully regulates WBAM while allowing variance in lower-level, intersegmental coordination patterns.

In this study, we provided visual information about the desired and actual step lengths at each foot-strike throughout all trials, including the perturbation and recovery steps. Participants were encouraged to achieve the target step lengths for as many steps as possible, and therefore participants may have relied on this feedback during perturbation recovery to return to their pre-perturbation walking patterns faster than they otherwise would without visual feedback. However, participants’ reactive response is unlikely to influence measures of momentum until late into the first recovery step as the step length information was only shown after the foot-strike of the first recovery step. It remains to be seen if patterns of interlimb coordination would differ in the presence of asymmetries that are not guided by online visual feedback.

Although the reactive intersegmental coordination patterns were significantly different from those observed during unperturbed locomotion, the overall patterns were qualitatively similar across steps. Taken together, these results may reflect two keys aspects of coordination during perturbed walking. First, the qualitative similarity between pre- and post-perturbation patterns observed here and in previous work [21] may reflect the dominant coordination patterns that characterize both unperturbed and perturbed bipedal walking. In contrast, the statistical differences between pre- and post-perturbation coordination patterns may reflect the changes in coordination necessary to maintain balance in response to perturbations. Patterns of intersegmental coordination observed during responses to external perturbations during walking likely capture a combination of passive limb dynamics, stereotypical pattern generation, and reactive balance control responses [37].

We observed that the upper limbs’ contribution to the control of angular momentum in the sagittal plane was negligible compared with lower limb segments during perturbation recovery. Since a stepping response is sufficient to restore balance from the treadmill accelerations used in this study, increases in momentum from the lower extremities may have been sufficient to restore sagittal plane WBAM. Consistent with our findings, Pijnappels et al. also found that arm movements had a small effect on body rotation in the sagittal plane during tripping over obstacles which elicits excessive forward rotation similar to the current study [38]. However, during larger perturbations that trigger backward falls, the arms elevate to shift the body’s center of mass back within the base of support [4]. This difference in the role of the arms across studies of perturbation recovery may result from the use of a larger velocity and displacement of the foot in the Marigold et al. [4] study. However, it remains to be seen how systematic variation of the magnitude and direction of external perturbations influences the role of the upper extremities during balance recovery.

Our results may also have implications for understanding the potential effects of interventions designed to reduce gait asymmetries in people post-stroke, as this is a common rehabilitation objective in this population [39]. Based on the current results, we would expect that reducing asymmetry in people post-stroke would also affect their reactive control strategies. However, further investigation is necessary to determine if reductions in asymmetry affect interlimb coordination during reactions to perturbations. The data from the current study illustrate how the intact neuromotor system modulates coordination between the upper and lower extremities in response to changes in asymmetry, and these data could serve as useful reference data to understand how sensorimotor impairments such as muscle weakness [40] and transmission delays [41] affect the ability to restore WBAM during perturbation recovery in people post-stroke.

## 5 Acknowledgments

We thank Natalia Sanchez, Ph.D., for her insights during the design of this experiment and Aram Kim for her assistance with the statistical analysis.

## 6 Author Contributions

C.L designed the experiment, collected data, analyzed data, and wrote the manuscript. J.M.F conceived of the experiment, advised in data analyses, and edited the manuscript.

## References

1. Winter DA. Human balance and posture control during standing and walking. Gait Posture. 1995;3: 193–214. doi:10.1016/0966-6362(96)82849-9

2. Tang PF, Woollacott MH, Chong RKY. Control of reactive balance adjustments in perturbed human walking: Roles of proximal and distal postural muscle activity. Exp Brain Res. 1998;119: 141–152. doi:10.1007/s002210050327

3. Winter DA. Biomechanics and Motor Control of Human Movement. Motor Control. 2009. doi:10.1002/9780470549148

4. Marigold DS, Bethune AJ, Patla AE. Role of the unperturbed limb and arms in the reactive recovery response to an unexpected slip during locomotion. J Neurophysiol. 2003;89: 1727–1737. doi:10.1152/jn.00683.2002

5. Wang T-Y, Bhatt T, Yang F, Pai Y-C. Adaptive control reduces trip-induced forward gait instability among young adults. J Biomech. 2012/02/28. 2012;45: 1169–1175. doi:10.1016/j.jbiomech.2012.02.001

6. Martelli D, Luciani LB, Micera S. Angular Momentum During Unexpected Multidirectional Perturbations Delivered WhileWalking. IEEE Trans Biomed Eng. 2013;60: 1785–1795.

7. Liu C, De Macedo L, Finley MJ. Conservation of Reactive Stabilization Strategies in the Presence of Step Length Asymmetries during Walking. Front Hum Neurosci. 2018;12: 251. doi: doi:10.3389/fnhum.2018.00251

8. Pijnappels M, Bobbert MF, Van Dieën JH. Push-off reactions in recovery after tripping discriminate young subjects, older non-fallers and older fallers. Gait Posture. 2005;21: 388–394. doi:10.1016/j.gaitpost.2004.04.009

9. Herr HM, Popovic M. Angular momentum in human walking. J Exp Biol. 2008;211: 467–81. doi:10.1242/jeb.008573

10. Popovic M, Hofmann A, Herr H. Angular momentum regulation during human walking: biomechanics and control. Robotics and Automation, 2004 Proceedings ICRA’04 2004 IEEE International Conference on. IEEE; 2004. pp. 2405–2411.

11. Simoneau GC, Krebs DE. Whole-Body Momentum during Gait: A Preliminary Study of Non-Fallers and Frequent Fallers. J Appl Biomech. 2000;16: 1–13. doi:10.1123/jab.16.1.1

12. Pijnappels M, Bobbert MF, Van Dieën JH. Contribution of the support limb in control of angular momentum after tripping. J Biomech. 2004;37: 1811–1818. doi:10.1016/j.jbiomech.2004.02.038

13. Vistamehr A, Kautz SA, Bowden MG, Neptune RR. Correlations between measures of dynamic balance in individuals with post-stroke hemiparesis. J Biomech. 2016;49: 396–400. doi:10.1016/j.jbiomech.2015.12.047

14. Nott CR, Neptune RR, Kautz SA. Relationships between frontal-plane angular momentum and clinical balance measures during post-stroke hemiparetic walking. Gait Posture. 2014;39: 129–34. doi: doi:10.1016/j.gaitpost.2013.06.008. Relationships

15. Silverman AK, Neptune RR. Differences in whole-body angular momentum between below-knee amputees and non-amputees across walking speeds. J Biomech. 2011;44: 379–385. doi:10.1016/j.jbiomech.2010.10.027

16. Honda K, Sekiguchi Y, Muraki T, Izumi SI. The differences in sagittal plane whole-body angular momentum during gait between patients with hemiparesis and healthy people. J Biomech. 2019;86: 204–209. doi:10.1016/j.jbiomech.2019.02.012

17. Lewek MD, Bradley CE, Wutzke CJ, Zinder SM. The Relationship Between Spatiotemporai Gait Asymmetry and Balance in Individuals With Chronic Stroke. J Appl Biomech. 2014;30: 31–36. doi:10.1123/jab.2012-0208

18. Latash ML, Ansen GJ, Anson JG. Synergies in Health and Disease: Relations to Adaptive Changes in Motor Coordination. Phys Ther. 2006;86: 1151–1160.

19. Aprigliano F, Martelli D, Tropea P, Pasquini G, Micera S, Monaco V. Ageing does not affect the intralimb coordination elicited by slip-like perturbation of different intensities. J Neurophysiol. 2017; jn.00844.2016. doi:10.1152/jn.00844.2016

20. Martelli D, Vashista V, Micera S, Agrawal SK. Direction-Dependent Adaptation of Dynamic Gait Stability Following Waist-Pull Perturbations. IEEE Trans Neural Syst Rehabil Eng. 2016;24: 1304–1313. doi:10.1109/TNSRE.2015.2500100

21. Aprigliano F, Martelli D, Micera S, Monaco V. Intersegmental coordination elicited by unexpected multidirectional slipping-like perturbations resembles that adopted during steady locomotion. J Neurophysiol. 2016;115: 728–740. doi:10.1152/jn.00327.2015

22. Plotnik M, Azrad T, Bondi M, Bahat Y, Gimmon Y, Zeilig G, et al. Self-selected gait speed - Over ground versus self-paced treadmill walking, a solution for a paradox. J Neuroeng Rehabil. 2015;12. doi:10.1186/s12984-015-0002-z

23. Sánchez N, Park S, Finley JM. Evidence of Energetic Optimization during Adaptation Differs for Metabolic, Mechanical, and Perceptual Estimates of Energetic Cost. Sci Rep. 2017;7: 7682. doi:10.1038/s41598-017-08147-y

24. Finley JM, Bastian AJ, Gottschall JS. Learning to be economical: the energy cost of walking tracks motor adaptation. J Physiol Physiol. 2013;591: 1081–1095. doi:10.1113/jphysiol.2012.245506

25. Reisman DS, Block HJ, Bastian AJ. Interlimb coordination during locomotion: What can be adapted and stored? J Neurophysiol. 2005;94: 2403–2415. doi:10.1152/jn.00089.2005

26. Song J, Sigward S, Fisher B, Salem GJ. Altered dynamic postural control during step turning in persons with early-stage Parkinson’s disease. Parkinsons Dis. 2012;2012: 386962. doi:10.1155/2012/386962

27. Havens KL, Mukherjee T, Finley JM. Analysis of biases in dynamic margins of stability introduced by the use of simplified center of mass estimates during walking and turning. Gait Posture. 2018;59: 162–167. doi:10.1016/j.gaitpost.2017.10.002

28. Kurz MJ, Arpin DJ, Corr B. Differences in the dynamic gait stability of children with cerebral palsy and typically developing children. Gait Posture. 2012;36: 600–604. doi:10.1016/j.gaitpost.2012.05.029

29. Reisman DS, Wityk R, Silver K, Bastian AJ. Split-Belt Treadmill Adaptation Transfers to Overground Walking in Persons Poststroke. Neurorehabil Neural Repair. 2009;23: 735–744.

30. Dempster WT. Space requirements of the seated operator: geometrical, kinematic, and mechanical aspects other body with special reference to the limbs. WADC Technical Report. 1955. pp. 1–254. doi: AD 087892

31. Hanavan EP. A Mathematical Model of the human body. Aerosp Med Rsearch Lab. 1964; 1–149.

32. Valero-Cuevas FJ, Klamroth-Marganska V, Winstein CJ, Riener R. Robot-assisted and conventional therapies produce distinct rehabilitative trends in stroke survivors. J Neuroeng Rehabil. 2016;13: 1–10. doi:10.1186/s12984-016-0199-5

33. Hothorn T, Bretz F, Westfall P. Simultaneous inference in general parametric models. Biom J. 2008;50: 346–363. doi:10.1002/bimj.200810425

34. Kuznetsova A, Brockhoff PB, Christensen RHB. lmerTest Package: Tests in Linear Mixed Effects Models. J Stat Softw. 2017;82. doi:10.18637/jss.v082.i13

35. Domkin D, Laczko J, Jaric S, Johansson H, Latash ML. Structure of joint variability in bimanual pointing tasks. Exp Brain Res. 2002;143: 11–23. doi:10.1007/s00221-001-0944-1

36. Papi E, Rowe PJ, Pomeroy VM. Analysis of gait within the uncontrolled manifold hypothesis: Stabilisation of the centre of mass during gait. J Biomech. 2015;48: 324–331. doi:10.1016/j.jbiomech.2014.11.024

37. Cappellini G, Ivanenko YP, Poppele R., Lacquaniti F. Motor Patterns in Human Walking and Running. J Neurophysiol. 2006;95: 3426–3437. doi:10.1152/jn.00081.2006

38. Pijnappels M, Kingma I, Wezenberg D, Reurink G, van Dieën JH. Armed against falls: the contribution of arm movements to balance recovery after tripping. Exp Brain Res. 2010;201: 689–699. doi:10.1007/s00221-009-2088-7

39. Patterson KK, Mansfield A, Biasin L, Brunton K, Inness EL, McIlroy WE. Longitudinal changes in poststroke spatiotemporal gait asymmetry over inpatient rehabilitation. Neurorehabil Neural Repair. 2015;29: 153–162. doi:10.1177/1545968314533614

40. Gray V, Rice CL, Garland SJ. Factors that influence muscle weakness following stroke and their clinical implications: a critical review. Physiother Canada. 2012;64: 415–26. doi:10.3138/ptc.2011-03

41. Sharafi B, Hoffmann G, Tan AQ, Y. Dhaher Y. Evidence of impaired neuromuscular responses in the support leg to a destabilizing swing phase perturbation in hemiparetic gait. Exp Brain Res. 2016;234: 3497–3508. doi:10.1007/s00221-016-4743-0

